# Evolutionary dynamics of polymorphic endogenous retrovirus insertions across wild house mouse populations

**DOI:** 10.1101/2025.09.23.678169

**Authors:** Tomoki Yano, Toyoyuki Takada, Kazumichi Fujiwara, Dai Watabe, Shutaro Hirose, Hiroshi Masuya, Toshinori Endo, Naoki Osada

## Abstract

Endogenous retroviruses (ERVs) represent a major source of structural variation in mammalian genomes, yet their diversity in wild populations remains poorly understood. Here, we conduct a comprehensive genome-wide survey of non-reference ERV insertions in wild house mice (*Mus musculus*) to characterize their distribution and evolutionary dynamics. Using a newly developed bioinformatics pipeline, we detected and annotated over 100,000 non-reference ERV insertions from short-read sequencing data across 163 wild mouse genomes.

Our analyses revealed marked differences in ERV insertion patterns among subspecies and populations, including variation in genomic localization and population-specific polymorphisms. These heterogeneous patterns suggest distinct evolutionary histories and host–retrovirus interactions across populations. For instance, we describe the distribution of the ERV-derived *Fv4* locus, which shows subspecies-restricted occurrence and confers resistance to murine leukemia viruses (MLVs). Several lines of evidence showed that the spread of *Fv4* insertions in Korean population has been driven by adaptive introgression from neighboring populations.

Our study provides the first large-scale population genomic scan of ERV diversity in wild house mice. By cataloguing extensive polymorphism in non-reference ERV insertions, our results highlight the role of ERVs as dynamic genomic elements that contribute to structural variation and adaptive evolution.

**Article Summary:** Endogenous retroviruses (ERVs) are viral sequences embedded in animal genomes that can create structural genetic variation. In this study, we conducted a genome-wide survey of non-reference ERV insertions in 163 wild house mice using short-read sequencing data and a newly developed computational pipeline. We identified more than 100,000 polymorphic ERV insertions and found substantial differences among subspecies and geographic populations. One example, the ERV-derived *Fv4* locus, illustrates how ERV variation can influence the genetic pattern of polymorphisms in the species. These results demonstrate that ERVs are dynamic genomic elements that contribute to population divergence and adaptive evolution.

## Introduction

Endogenous retroviruses (ERVs) are transposable elements derived from RNA viruses that encode reverse transcriptase and become endogenized in host genomes through integration into germline DNA. ERVs are classified into three major classes (Class I–III) (Stoye and Coffin 1987; Stocking and Kozak 2008) and collectively comprise approximately 5% of both human and house mouse reference genomes (Lander, et al. 2001; Mouse Genome Sequencing, et al. 2002; Nellåker, et al. 2012). ERVs can alter organismal phenotypes through multiple mechanisms, including gene silencing, insertional disruption, and regulatory activation via long terminal repeats (LTRs). In addition to these functions, several ERV or ERV-like elements have been shown to influence resistance to retroviral infection (Ikeda, et al. 1981; O’Brien, et al. 1983; Best, et al. 1996; Dodding Mark, et al. 2005; Wu, et al. 2005; Zhang, et al. 2008), and studies of wild house mice suggest that the distribution of ERV-derived restriction factors may reflect historical adaptation to viral pathogens (Boso, et al. 2021; Fujiwara, et al. 2024). Similar antiviral functions of ERVs have been raised in humans (Frank, et al. 2022), yet direct evidence remains scarce. These observations highlight the need for comprehensive, population-scale surveys of ERV insertions in natural populations to understand the mechanisms by which the phenotypic variation has been shaped by originally exogenous genetic components.

In laboratory house mice (*Mus musculus*), including both classical and wild-derived inbred strains, extensive variation in ERV insertion patterns has been documented (Nellåker, et al. 2012; Ferraj, et al. 2023), yet the genetic diversity of ERV polymorphisms in wild populations remains largely unexplored. Here, we focus on non-reference polymorphic ERV insertions segregating in wild house mice. House mice comprise three major subspecies, *M. m. castaneus*, *M. m. domesticus*, and *M. m. musculus* (Boursot, et al. 1996; Fujiwara, et al. 2022), whereas classical inbred strains predominantly possess a *domesticus*-like genetic background (Keane, et al. 2011). Because the reference genome derives from C57BL/6J (Mouse Genome Sequencing, et al. 2002), it lacks many ERV insertions present in natural populations. Given the extensive genetic diversity among wild mice and the presence of multiple active ERV families, characterizing non-reference insertions provides insight into the population genetic processes shaping ERV variation. Moreover, house mice remain susceptible to active murine leukemia viruses (MLVs), particularly class I retroviruses such as Friend and Moloney leukemia viruses. The interplay between exogenous retroviruses and resident ERVs may therefore leave detectable evolutionary signatures in wild genomes.

However, systematic detection of ERV insertions in non-human species has been hindered by both technological and analytical constraints. Long-read sequencing platforms such as Oxford Nanopore and PacBio can resolve complex structural variants (Logsdon, et al. 2020; Chu, et al. 2021), but sequencing large numbers of individuals with these technologies remains prohibitively expensive. In contrast, short-read whole-genome datasets are widely available across species, yet existing ERV detection tools were largely designed for human applications, often exhibit elevated false-discovery rates, and scale poorly with large sample sizes (Keane, et al. 2012; Gardner, et al. 2017; Chen and Li 2019; Bowles, et al. 2022; Kojima, et al. 2023). Searching for a broad spectrum of transposable elements also incurs substantial computational cost, limiting practical application in population genomic studies. To address these limitations, we developed ERVscanner, a new ERV detection pipeline optimized for high computational speed and low false-discovery rates. We benchmarked ERVscanner using the wild-derived strain JF1 (Koide, et al. 1998; Takada, et al. 2013; Takada, et al. 2025) with both short- and long-read sequencing data, confirming accurate recovery of known insertions.

To demonstrate the utility of ERVscanner and to explore potential adaptive ERV insertions, we examined *Fv4*, a well-studied ERV-derived restriction factor absent from the reference genome. *Fv4* encodes an env protein derived from a class I retrovirus (Odaka, et al. 1978; O’Brien, et al. 1983; Inaguma, et al. 1991), exhibits presence/absence polymorphism, and is most common in *M. m. castaneus*. In East Asia, secondary contact between subspecies *castaneus* and *musculus* forms a broad hybrid zone (Fujiwara, et al. 2022), which can facilitate introgression of potentially adaptive alleles (Huerta-Sanchez, et al. 2014). Although Korean house mice display predominantly *musculus*-like genomic backgrounds, many individuals harbor *Fv4* (Inaguma, et al. 1991; Boso, et al. 2021). Our analyses indicate that *Fv4* was recently introgressed from the subspecies *castaneus* into *musculus* and that surrounding genomic patterns are consistent with a selective sweep, supporting previous functional evidence for its role in retroviral resistance.

Application of the same framework revealed three additional ERV insertions showing population patterns compatible with introgression in Korean mice. Although their functional relevance remains unclear, these loci represent promising candidates for further investigation and illustrate how scalable ERV discovery can reveal previously unrecognized variation. Overall, our study provides a practical tool for ERV detection in short-read datasets, a population-scale catalogue of ERV insertions in wild house mice, and prioritized loci, including *Fv4*, for future functional and ecological analysis.

## Materials and Methods

### Algorithm of ERVscanner

ERVscanner is a pipeline designed to detect non-reference insertions of ERV sequences using short-read mapping files (BAM or CRAM) as input. All codes are distributed at Github (https://github.com/Qtom-99/ERVscanner). Genotypes of samples are output as a single VCF file. It consists of four main processes: read filtering, identification of read clusters around insertions, filtering based on insertion content, and genotyping. A graphical representation is provided in Supplementary Figure 1 and 2. Although ERVscanner is specifically designed for ERV sequences, it can be applied to other classes of mobile elements. The pipeline utilizes pre-defined ERV regions in the reference genome, as annotated by the Dfam database (Storer, et al. 2021), which identifies these regions using a hidden Markov model. We refer to these annotated regions as masked regions (MRs). For further processing, ERVscanner requires BED-format files containing MRs and FASTA-format sequences of MRs. The MR sequences are reformatted to the sense (+) strand representation of ERVs, and consensus sequences of ERVs are included in the FASTA file in the preprocess step. Additionally, any missing ERV sequences that are not represented in the reference genome can be manually added to the FASTA file.

Using the BED file, ERVscanner first extracts discordantly mapped read pairs after removing PCR duplicates. Discordant pairs include those mapped to different scaffolds/chromosomes, separated by more than 10 kb, or mapped with incongruent orientations.

The extracted pairs are filtered based on MRs. When one read maps within an MR and the mate maps outside an MR, only the read mapped outside the MR is retained if it has high mapping quality (MQ ≥ 30). Read pairs in which both reads map within MRs are excluded from further analysis. Because detecting insertions within MRs is challenging with short-read data and prone to elevated false discovery rates, ERVscanner estimates insertions only from reads supported by high-confidence mappings outside MRs.

If reads were aligned using BWA-MEM or other ALT-aware mappers against non–telomere-to-telomere reference genomes, reads mapping to both primary and alternative/unplaced contigs may receive artificially high mapping quality scores on primary chromosomes. To reduce false positives, clusters composed of reads with secondary alignments to alternative/unplaced contigs were removed.

After filtering, ERVscanner identifies clusters indicative of non-reference ERV insertions. Insertions are expected to generate forward–reverse read clusters flanking the breakpoint. For each orientation, regions covered by reads separated by ≤200 bp are merged. Clusters exceeding a predefined read-count threshold are defined as forward or reverse clusters. When forward and reverse clusters occur within 200 bp of each other with the expected orientation, the locus is classified as a candidate non-reference insertion site.

For each locus, we estimated the insertion content by re-mapping reads paired with the cluster-supporting reads, using BWA against 361 non-redundant transposable element sequences in the genus *Mus* from Dfam (Supplementary Table 3). We further reassigned Long_Terminal_Repeat families to ERV1 and MaLR families to ERV3 according to previous studies (Stocking and Kozak 2008; Kawase and Ichiyanagi 2023). Hereafter we refer to these library sequences as the Dfam NRPH (non-redundant profile HMM) consensus. The Dfam NRPH consensus represents the consensus sequences of transposable elements without overlapped annotations.

To improve accuracy, we aggregated short-read sequences from multiple individuals under the assumption that polymorphic insertions at the same locus share identical insertion contents. If the insertion sequence is correctly inferred, reads from the left and clusters are expected to map in consistent orientations (Supplementary Figure 2). For each mapped library sequence, we counted the number of left- and right-flanking reads mapped, and retained the smaller value as the support score. After calculating support scores for all candidate library sequences, we assigned to the sequence with the highest support score. Candidate loci with a support score ≥3 were defined as valid clusters.

For each valid cluster, we genotyped the insertions to determine whether insertions were heterozygous or homozygous. Within each valid cluster, clipped reads were extracted using CIGAR information to identify precise breakpoints. If clipped reads from forward and reverse clusters overlapped around a breakpoint, we assigned the minimum position as the breakpoint, in order to maintain consistency with the VCF format generated by general genotyping software. Heterozygous insertions were identified when at least three breakpoint-crossing reads covering a 30-bp window centered on the breakpoint. Valid clusters lacking a sufficient number of clipped reads were genotyped as ‘1/.’ in the output VCF file, indicating the presence of an insertion but uncertainty regarding zygosity.

### Evaluation of insertion calling using long-read sequences

Variant data based on long-read sequencing of the JF1 strain were downloaded from a publicly available website (https://molossinus.brc.riken.jp/pub/MoGplus30data/). The short-read data (DRA006630) was randomly down sampled to 30x coverage. Since the variant data was based on GRCm39, the coordinates of variants were converted to GRCm38 using the liftOver tool. To infer insertion contents from long-read sequencing data, we extracted insertion sequences ranging from 200 to 10,000 bp and performed a homology search using BLASTN (Altschul, et al. 1990) against Dfam NRPH consensus sequences. For each insertion, we identified the longest hit with E-value ≥30 and ≥50% query coverage to classify the insertions by class and family. Insertions assigned to the ERV classes were selected for further analysis.

ERVcaller 1.4 was run using Dfam NRPH consensus sequences as seed sequences for insertions (Chen and Li 2019). Because we were unable to complete the analysis of the JF1 strain using the default ERVcaller parameters, we reduced the internal parameter specified by -c in BWA-MEM (Li and Durbin 2009) for read extraction to 1/10 of the original value (from 100,000 to 10,000). This modification significantly reduced the running time but may have affected accuracy to some extent. After running ERVcaller, we extracted insertions classified as the ERV classes. The output VCF file is accessible at Zenodo (DOI: 10.5281/zenodo.18695322).

ERVscanner was used to estimate ERV insertions as described above. We first merged adjacent insertions within 40 bp within each dataset, and then considered insertions from different datasets to be overlapping when their sites were within 40 bp.

### Analysis of 163 house mouse short-read data

Short-read alignment files of 163 wild *M. musculus* and 7 *M. spretus* samples determined by Fujiwara et al. (2024) were used for the analysis. Briefly, the file was constructed high-coverage whole-genome short-read sequences by mapping to the GRCm38 reference genome.

We downloaded the annotation of ERV sequences for GRCm38 from Dfam database. Segments classified as ERV1 and ERV2-group were considered as candidate ERV sequences. The ERV2-group class includes ERV2, 3, 4, and Lentivirus families. In addition, RLTR4_MM-int family, which is not classified as the ERV class but classified as the Long_Terminal_Repeat_Element class, was included in the analysis because previous study classified RLTR4_MM-int as MLV-related sequences (Kawase and Ichiyanagi 2023). Using the information, we constructed BED and FASTA files to represent the MRs. To further characterize known ERV-derived restriction factors, we added the sequences of *Fv4*, *RMCF1*, and *RMCF2* in the public database (DDBJ/EMBL/Genbank accession numbers: AF490353, AF490352, AY999005, AH001845, M33884, and AH001894)

We inferred insertions sites using ERVscanner, explained above, using five reads for the thresholds for each forward- and reverse-cluster, and three reads for the thresholds of breakpoint-crossing reads to judge the zygosity. Principal Component Analysis (PCA) was performed using scikit-learn library of Python.

### Detection of selective sweep

Population genetics summary statistics were calculated to detect the signature of selective sweep surrounding the non-reference ERV insertions. Nucleotide diversity (*π*), Tajima’s *D, and F*_ST_ between Korean (11 individuals from Korean peninsula) and Northeastern Chinese (five individuals from Liaoning, Jiling, and Heilongjiang Provinces) samples were calculated using VCFtools (Danecek, et al. 2011), for 5-kbp-lengh windows with step size of 1000 bp.

For the calculation of nSL and XP-nSL statistics, we assigned ancestral alleles using the genotypes of seven *M. spretus*, and modified the VCF file using the information. Only the monomorphic sites in *M. spretus* were used for the analysis. Both statistics were calculated using selscan 2.0 software with default parameters (Szpiech 2021). For XP-nSL, we used Korean samples for the test population and Northeastern Chinese samples for the reference population.

Genealogies surrounding the insertion sites of *Fv4* were inferred using RELATE software, using the same process described in Fujiwara et al. (2024), but in this study, we re-inferred population size change using only the chromosome 12 sequences of subspecies *musculus* samples with five rounds of iterations. In total 100 branch lengths were resampled using RELATE for CLUES.

### Candidate of ERV insertions under adaptive evolution

We selected SNPs from Korean populations located within regions under selective sweep, identified using nSL statistics. SNPs in the top 0.1% of nSL scores were then pruned using PLINK with the --indep-pairwise 20 5 0.8 option. After pruning, we selected SNPs that (1) lacked exons within 10 kb and (2) were located within 1 kb of ERV insertions. Loci were retained if the allele frequency of the ERV insertion was equal to or greater than 0.5 in the Korean population and higher than that in the subspecies *castaneus*.

## Results

### Identification of ERV insertions in JF1 mouse using a long-read sequencer

We first characterized non-reference ERV insertions in the JF1 mouse strain, identified using high-depth long-read genome sequence data (Takada, et al. 2025). In total, we identified 6558 insertions and 810 deletions, ranging from 200–10,000 bp and homologous to ERV sequences. As observed in previous studies, the most abundant class of insertions was Class II (ERV2/ERVK), including IAP elements, comprising 77.2% of all ERV insertions. The numbers of insertions for Class I (ERV1) and III (ERV3/ERVL and ERVL-MaLR) were nearly equivalent: 10.8% for Class I and 12.0% for Class III. The number of deletions showed a similar trend. Among Class I, the MuLV family, including internal sequences (MuLV-int), was the most abundant.

The distributions of insertion/deletion length are shown in Supplementary Figure 3. While the majority of insertions and deletions are short, including fragmented copies and solo LTRs, 2236 insertions and 126 deletions were in the range between 2000 and 10,000 bp..

### Evaluation of ERV insertion detection using short-read data

In order to effectively handle large-scale population genomic data of short-read genome sequences, a new bioinformatics pipeline, ERVscanner, was developed. The basic idea of ERVscanner is almost the same as the previously published programs for detecting mobile elements, but ERVscanner largely relies on the repetitive sequence annotation for the reference genome by Dfam database (Storer, et al. 2021). Assuming that nearly all non-reference ERV insertions have homologous sequences annotated as ERV regions in the reference genome, we can significantly reduce the computational search space. Additionally, ERVscanner avoids assembling insertion sequences, a process that is typically time-consuming, because such assemblies are often error-prone due to the nature of short-read sequences and high complexity of insertion sequences. Instead of inferring the exact sequence of insertions, ERVscanner estimates the class, family, and orientation of candidate insertions for each individual, outputting the results in VCF format. We can also incorporate other seed sequences for potential insertions. In this study, we used sequences of *Fv4*, *RMCF1*, and *RMCF2*, which are known antiviral factors derived from retroviruses, as additional targets for non-reference insertions. Detailed methods are described in the Methods section. Note that ERVscanner is not designed to investigate deletions, as identifying deletions of repetitive sequences using short-read sequencing is highly challenging.

We evaluated the computational speed, sensitivity, and accuracy of ERVscanner using high-coverage long-read and short-read sequences from the JF1 mouse strain, assuming that long-read data accurately represent true insertion sequences. Previous studies estimated the false-discovery rate (FDR) of insertions using long-read sequencing was around 5.8%, examined by PCR amplification (Ferraj, et al. 2023). ERVscanner is designed to enhance accuracy by leveraging data from multiple individuals. However, for benchmarking purposes, we evaluated the performance using only a single individual of the JF1 strain to ensure a fair comparison. In addition, the sequence depth was limited to 30-fold by downsampling the original data. For comparison, we selected ERVcaller, a tool noted for its high sensitivity and low false-positive rate in a benchmarking study (Bowles, et al. 2022). Both pipelines analyzed all classes of ERVs using reference-mapped short-read files. Utilizing eight cores of an Intel Xeon CPU E5-2620 v4 at 2.10 GHz, ERVscanner detected 6177 insertions in approximately 3 hours. For ERVcaller, we estimated all non-redundant transposable elements rather than only ERV sequences and extracted inferred insertions matching ERVs to enhance accuracy. Using stringent criteria (support levels 4 and 5), ERVcaller identified 11,850 candidate ERV insertions in 82 hours with the same computer used for ERVscanner.

Figure 1 presents a Venn diagram comparing the results of JF1 strain using three datasets. As expected, many insertions identified by long-read sequencing are undetectable by short-read methods, showing the inherent limitations of short reads in detecting ERV insertions. ERVcaller identified more correct ERV insertions than ERVscanner; however, its FDR approached 59%. In contrast, ERVscanner exhibited an FDR of approximately 28%. While this rate remains high, it is notably lower than that of ERVcaller. The majority of mispredictions in ERVscanner are due to misclassification of the sequence class rather than incorrect insertion detection (data not shown). Notably, ERVscanner successfully identified the insertion of an endogenous retrovirus *β*4 (ERVB4) element in the *Ednrb* gene, responsible for the characteristic piebald coat color phenotype of JF1 mice (Yamada, et al. 2006; Tanave and Koide 2020).

**Figure 1.**
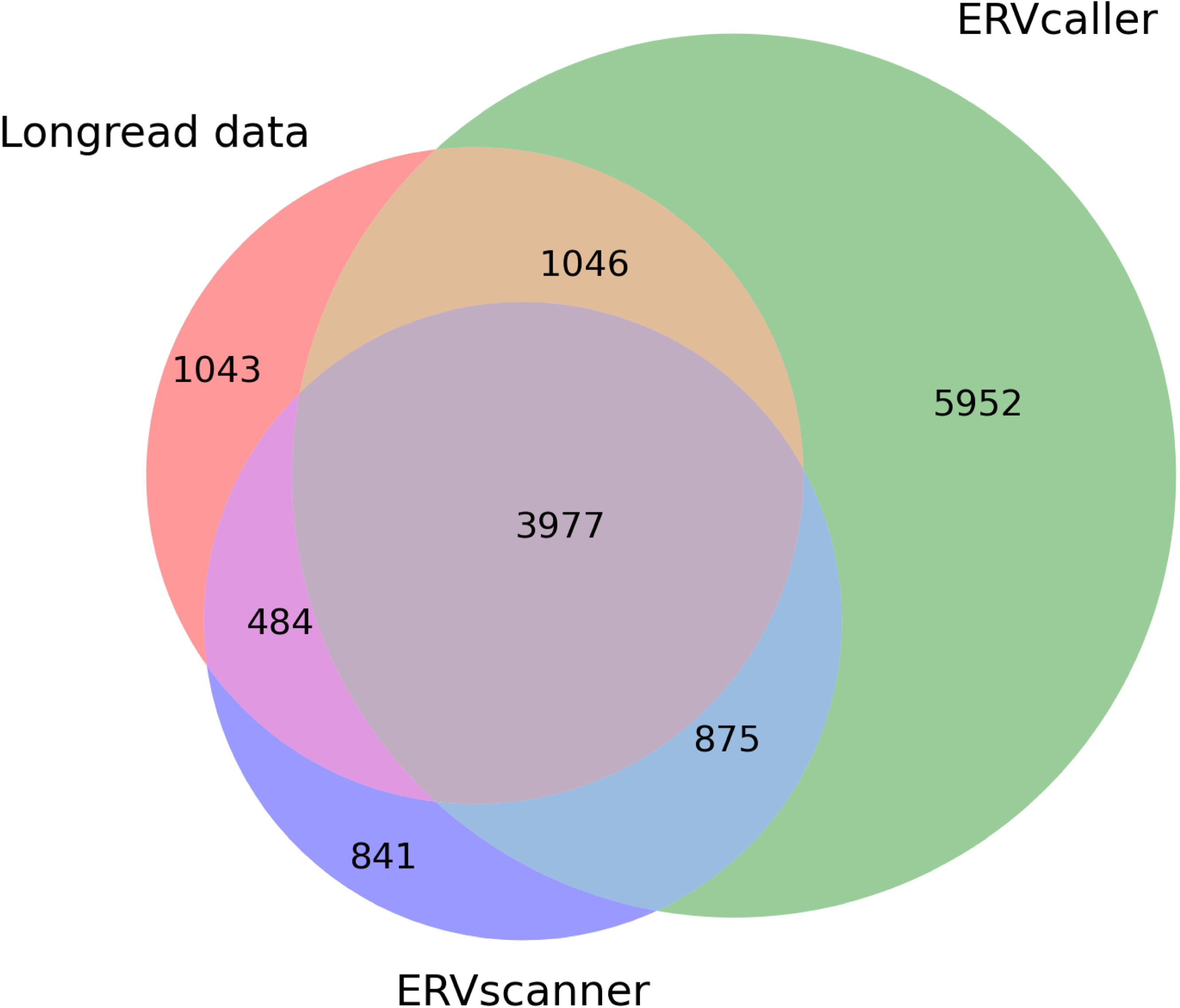
Venn diagram of detected non-reference ERV insertions in the JF1 strain genome. Long-read represents high-coverage PacBio data, while ERVcaller and ERVscanner utilized 30x short-read data.

### Population genomic search of ERV across wild house mouse genomes

We analyzed short-read mapping data from 163 wild house mice using ERVscanner (Supplementary Table 1), and identified an average of 7103, 3524, and 6996 non-reference ERV insertions on autosomes per individual in the subspecies *castaneus*, *domesticus*, and *musculus*, respectively. The distribution of these insertions for each individual is presented in Figure 2. As expected, the subspecies *domesticus* harbored the fewest ERV insertions, because the reference genome is primarily derived from the subspecies *domesticus*. We showed that the class composition of ERV insertions differs among subspecies, with significant differences detected for each ERV class (Kruskal–Wallis test; Class I: *p* = 1.21 × 10^−24^; Class II: *p* = 9.53 × 10^−15^; Class III: *p* = 3.54 × 10^−23^). Further pairwise tests suggested that ssp. *musculus* tends to harbors more Class I and II insertions and fewer Class III insertions, compared with ssp. *castaneus* and *domesticus* (all adjusted *p* < ×10^−7^; Dunn’s test after Holm correction).

**Figure 2.**
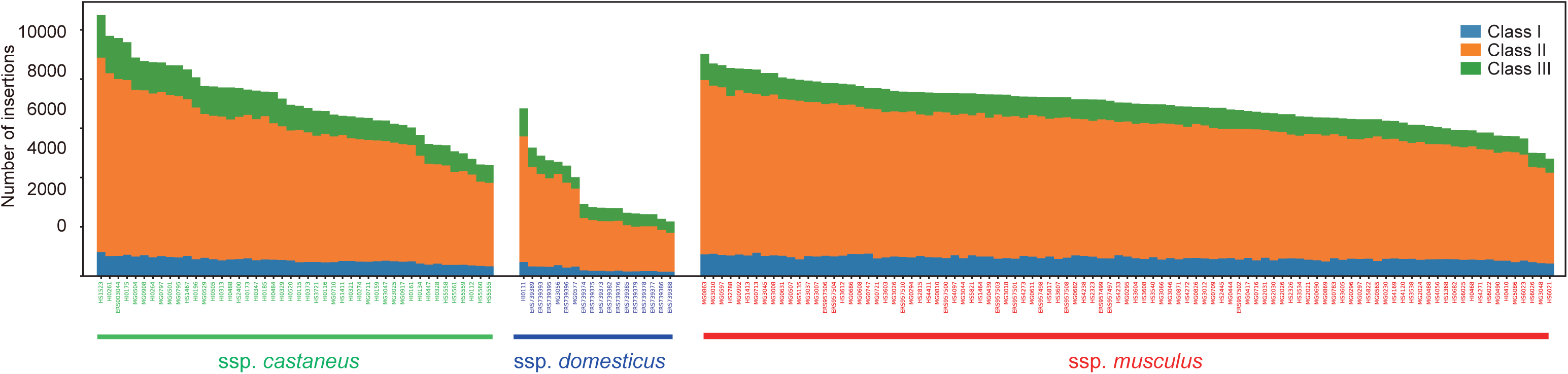
Histogram of loci with non-reference ERV insertions among all wild house mouse individuals. The labels of subspecies were determined by the majority of genomic components inferred by ADMIXTURE software (Fujiwara et al. 2024).

We asked whether global population structures could be reconstructed using the pattern of non-reference ERV insertions. PCA was performed using ERV insertions as markers, and the results reflected the subspecies population structures of wild house mice (Supplementary Figure 4).

### Functional effect of ERV insertions

In total, we identified 178,334 insertion loci that were not present in the reference genome. As previously reported, there was a strand bias of ERV insertions in intronic regions. We observed 62,720 insertions overlapping with the transcribed regions, and among them, 41,191 ERVs were inserted in a reverse orientation to the transcription.

We further investigated the relationship between integration sites and local recombination rates. We first calculated average local recombination rates and number of insertions for each 20kbp-length window and computed Spearman correlation coefficient, but did not observe a significant correlation (*ρ* = 0.0026, *p* = 0.45). We next investigated the association between insertion allele frequency and local recombination rate, and found a statistically significant but weakly negative correlation, suggesting that common insertions preferentially reside in low-recombination regions (Spearman correlation coefficient: −0.0158; *P* value: 7.20 × 10^−11^; Supplementary Figure 5).

We identified 320 genes with 325 ERV insertions disrupting protein-coding regions (Supplementary Table 2). Functional enrichment analysis using Metascape revealed significant enrichment of several gene clusters (Supplementary Figure 6). For example, genes associated with response to chemical stimulus (GO:0001580), including 64 olfactory receptor (OR) genes and 14 vomeronasal receptor (Vmnr) genes, were significantly enriched. Other overrepresented categories included metabolism of small nitrogenous compounds (mmu00410; WP522) and immune system–related processes (GO:0002218), such as macrophage migration, monocyte chemotaxis, and leukotriene metabolism.

### Polymorphic insertion of restriction factor Fv4

To examine the causal relationship between ERV insertions and natural selection, we focused on the *Fv4* gene, a major restriction factor for MLV infection that is absent from the current reference genome. Long-read sequencing data revealed that the JF1 mouse strain possesses *Fv4* at chr12:80,797,701 in the autosomal regions of GRCm38 reference genome. ERVscanner successfully inferred that *Fv4* had presence/absence polymorphism in wild house mouse samples at the same position. The geographic distribution of *Fv4* (Supplementary Figure 7) was consistent with the findings of Boso et al. (2021) but offers higher geographic resolution. In general, East Asian house mice exhibit a *musculus*-like genetic background in northern China and a *castaneus*-like background in southern China, with the Yangtze River (or Qinling Mountains) serving as an approximate boundary. *Fv4* is primarily restricted to the subspecies *castaneus*, but some East Asian *musculus*-like individuals along the eastern coastal regions, including northern China (Shandong and Hebei Provinces), Korea, Japan, and Russian Primorye, also harbor *Fv4*. Among these, Japanese and Russian samples show admixed subspecies origins. In contrast, seven individuals from northeastern China, which have a strongly *musculus*-like genetic background, do not harbor *Fv4*.

The distribution pattern of *Fv4* suggests that the gene may have introgressed from the subspecies *castaneus* to *musculus* during secondary contact in East Asia. We computed local *f*_4_ statistics using the configuration *f*_4_(SPR, S-CHN; KOR, NE-CHN), which becomes negative if introgression occurred from the southern Chinese population (S-CHN) into the Korean population (KOR), but not into the northeastern Chinese population (NE-CHN), with *M. spretus* (SPR) as the outgroup. As shown in Supplementary Figure 8, the *f*_4_ statistics were strongly skewed toward negative values around the *Fv4* insertion sites, indicating that the genomic segments surrounding the *Fv4* insertions were introgressed into the Korean population but not into the northeastern Chinese population.

To further investigate whether the spread of *Fv4* has been driven by adaptive evolution, we conducted a genomic scan for positive selection in Korean samples. We plotted nucleotide diversity and Tajima’s *D* statistics surrounding the *Fv4* insertion sites (Figure 3) using SNP data. Additionally, using northeastern Chinese samples, which do not possess *Fv4*, as a reference, we calculated and plotted *F*_ST_ and cross-population normalized selection scores (XP-nSL) (Figure 3). The results indicate a reduction in polymorphisms, an excess of rare alleles, high differentiation between populations, and homogeneous haplotype structures, all signatures consistent with a selective sweep. Notably, SNPs near the *Fv4* insertion sites fell within the lowest 0.01% of rankings among SNPs on the chromosome. These findings support the hypothesis that *Fv4* has undergone adaptive introgression from the subspecies *castaneus* to the subspecies *musculus* in East Asia, likely facilitated by secondary contact between these subspecies. The presence of *Fv4* in Korean *musculus*-like populations, despite their genetic background, suggests positive selection has favored the retention of this gene, potentially due to its role in conferring resistance to MLVs.

**Figure 3.**
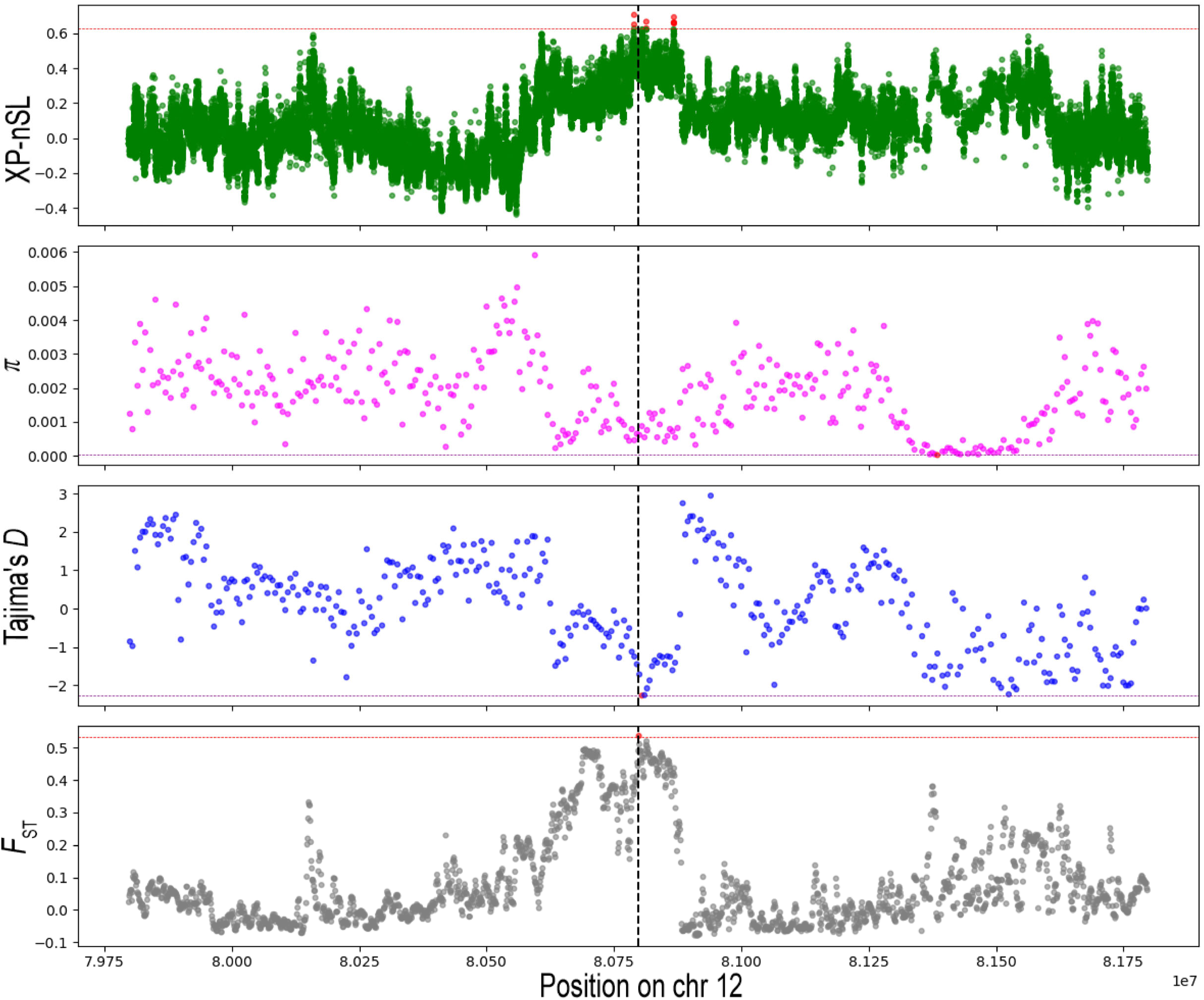
Genomic signature of selective sweep around the insertion of Fv4 in the Korean population. The vertical dashed line in the center of plot represents the estimated insertion site of Fv4. Dashed line in red and purple show the 0.01% thresholds for the top and bottom distributions, respectively. For the XP-nSL and F_ST_ calculation, northeastern Chinese individuals, which also possess strong M. m. musculus genetic background, were used as the control population.

We also inferred the timing and strength of selection using genealogy-based methods implemented in the RELATE and CLUES software (Speidel, et al. 2019; Stern, et al. 2019). We estimated that the increase of *Fv4* insertion frequency occurred after approximately 1000 generations ago, with a selection coefficient of 0.47%. Log-likelihood ratio against the neutral model was 8.50. The trajectory of *Fv4* insertions over time in the subspecies *musculus* is shown in Supplementary Figure 9.

### Potential candidates of drivers for adaptive ERV insertions

We conducted a genomic scan for selective sweeps in the Korean population using nSL statistics. We selected this population because it has a relatively homogeneous population structure and a sufficient sample size. From this analysis, we selected the top 0.1% of SNPs with the highest nSL scores, resulting in 607 high-nSL SNPs. Among these SNPs, 65 SNPs lacked adjacent annotated exons within 10 kbp but were flanked by ERV insertions within 1 kbp. Among the candidate insertions, three insertions present at high frequencies in the Korean population but low in the subspecies *castaneus*. In all three cases, the JF1 mouse strain possesses the corresponding insertions, and we inferred the sequence contents using available long-read sequencing data.

Table 1 presents a list of insertions that have potentially reached high frequency in the Korean population by adaptive evolution. Among these, one locus (chr1:3723618) is a solo LTR derived from an IAP element; another (chr7:42156902) is a nearly full-length RLTR10 insertion lacking fully encoded proteins; and a third (chr18:21435569) encodes a 1,153-amino-acid protein in the longest open reading frame, corresponding to ERV gag and pol proteins.

**Table 1.**

| Position | Class | Name | Length | Freq. Korea | Freq. *castaneus* |
| --- | --- | --- | --- | --- | --- |
| chr1:3723618 | ERV2 | IAPLTR1a_Mm | 764 | 0.89 | 0 |
| chr7:42156902 | ERV2 | RLTR10 | 2685 | 0.94 | 0.08 |
| chr18:21435569 | ERV1 | RLTR6B_Mm | 3387 | 0.76 | 0.17 |

## Discussions

### Evaluation of methods

In this study, we conducted a comprehensive search for non-reference ERV insertions in wild house mouse genomes and identified 178,334 potentially polymorphic insertions across 163 individuals. To facilitate this analysis, we developed a scalable pipeline designed to efficiently handle large datasets. Our pipeline is optimized for population genomic data and leverages shared insertions among individuals to enhance detection accuracy.

Although some insertions may result from somatic integration of exogenous retroviruses such as MLVs, 80,036 loci were shared among multiple individuals, suggesting that these insertions originated in germline lineages. Based on our benchmarking results, the false-positive and false-negative rates were nearly equivalent, suggesting that overestimation and underestimation largely offset each other. Therefore, the estimated total number of non-reference ERV insertions is unlikely to be substantially biased.

The total number of insertions identified here exceeds that reported in recent long-read sequencing studies based on a limited number of inbred strains. The observed difference is expected, as the genetic diversity represented by currently used inbred strains accounts for only a small fraction of the total diversity within the species. Overall, our results suggest that ERV insertions make a substantial contribution to the genetic diversity of wild house mice.

### Genomic pattern of ERV insertions

Multiple genomic factors may contribute to the distribution patterns of ERVs across genomes. ERVs and other transposable elements tend to integrate into regions of open chromatin, while recombination may facilitate their removal (Campos-Sánchez, et al. 2016) (Kent, et al. 2017). Additionally, some ERVs contain sites that promote recombination (Edelmann, et al. 1989), whereas ERV integration into functional genomic regions can negatively impact fitness and may be selectively removed from the genome (Campos-Sánchez, et al. 2016). These opposing forces make it difficult to predict whether ERVs preferentially found in high or low recombination rates.

Our data focusing on polymorphic ERVs did not reveal association between the density of ERV insertions and local recombination rate. In contrast, we observed a significant negative correlation between the allele frequency of ERV insertions and recombination rate, such that insertions tend to be at lower frequency in regions of high recombination.

This pattern is consistent with the possibility that purifying selection against ERV insertions is more efficient in high-recombination regions, where linkage effects are reduced. In addition, recombination-associated processes, such as deletion via non-allelic homologous recombination, may further decrease the persistence of ERV insertions in these regions. These mechanisms could lead to an excess of low-frequency insertions in high-recombination regions, even if the overall insertion density shows little association with recombination rate.

### Genes affected by ERV insertions

Several observations suggest that polymorphic ERV insertions may have functional consequences. The overrepresentation of OR genes likely reflects relaxed purifying selection acting on this large and partially pseudogenized gene family, highlighting how insertion tolerance varies across the genome. In addition, enrichment of genes involved in metabolic pathways raises the possibility that ERV insertions contribute to standing functional variation in wild populations. ERVs may also modify gene regulation. The ERVB4 insertion within *Ednrb*, which underlies the coat color phenotype of JF1 mice, illustrates how such events can have pronounced phenotypic effects.

### ERV insertions and adaptation

While ERV insertions can influence organismal phenotypes in various ways, one intriguing hypothesis is that ERVs may act as restriction factors against similar exogenous retroviruses (Goff 2004). However, molecular evidence for this mechanism remains scarce. In this study, we searched for retroviral-derived restriction factors not presented in the reference genome, *Fv4*, *RMCF1*, and *RMCF2*, but only *Fv4* was detected by our pipeline. *Fv4*, a restriction factor that enhances resistance to MLVs but is absent from the mouse reference genome, serves as an ideal model for studying the evolutionary dynamics of ERV insertions in mammalian immunity. Derived from MLVs, *Fv4* retains only the *env* gene in complete form, which encodes an envelope protein with several substitutions and blocks MLV receptors, thereby inhibiting infection. The gene likely originated in the common ancestor of the subspecies *castaneus*, but previous studies have identified *Fv4* in some individuals of the subspecies *musculus*.

Our global population-wide survey revealed that individuals from northern China, Korea, Japan, and Russian Primorye carry *Fv4*. Notably, the northern Chinese and Korean samples predominantly exhibit *M. m. musculus* genetic backgrounds, suggesting that *Fv4* spread in these populations despite restricted gene flow, likely due to its strong selective advantage. Population genetic analyses, using various summary statistics, detected strong selective sweeps around *Fv4* insertion sites, supporting a scenario of adaptive introgression from the subspecies *castaneus* to *musculus* in this region. Genealogical analysis with RELATE software estimates the spread occurred approximately 1000 generations ago. Notably, the signature of selection was detected using only SNP data. If we had not investigated the polymorphic insertions of *Fv4*, the driving force of strong selective sweep would have been invisible to us.

Previous study has also revealed the adaptive introgression of another well-studied restriction factor, *Fv1* in the Japanese population. *Fv1*, encoding gag protein derived from MuERV-L, is present in the reference genome, but has two distinct alleles with different protein length, B and N, showing restriction to different types of MLVs. The distribution of N and B alleles is characteristic among subspecies, and Fujiwara et al. (2024) showed that the *castaneous*-type B allele has been adaptively introgressed into the Japanese population. The examples of *Fv1* and *Fv4* indicate the adaptive introgression of ERV-derived restriction factors play a key role in the evolution of genomes under secondary contact of genetically differentiated populations. Although we lack direct experimental evidence for the functional and selective effects of the ERV insertions shown in Table 1, these candidates could serve as valuable resources for studying the relationship between ERV integrations and adaptive evolution.

## Conclusions

Taken together, the findings in this study highlight ERVs as a source of genetic diversity with the potential to influence host adaptation. By providing both a validated adaptive introgression event (*Fv4*) and a set of prioritized candidate loci, our study generates testable hypotheses regarding the evolutionary consequences of ERV insertions. More broadly, ERVscanner establishes a practical approach for detecting such polymorphisms in short-read datasets, enabling scalable discovery in systems where long-read sequencing remains limited. The catalogue and framework presented here offer a foundation for both comparative studies across mouse populations and extension to other species, promoting the integration of ERV dynamics into mainstream population and functional genomics.

## Data Availability Statement

All nucleotide sequence data used in this study were obtained from public databases. The accession numbers are listed in Supplementary Table 1. The ERVscanner code is available on GitHub (https://github.com/Qtom-99/ERVscanner). The list of ERV insertions among 163 wild house mice is available at Zenodo (DOI: 10.5281/zenodo.18695322) in VCF file format.

## References

Altschul SF, Gish W, Miller W, Myers EW, Lipman DJ. 1990. Basic local alignment search tool. Journal of Molecular Biology 215:403–410.

Best S, Tissier PL, Towers G, Stoye JP. 1996. Positional cloning of the mouse retrovirus restriction gene Fvl. Nature 382:826–829.

Boso G, Lam O, Bamunusinghe D, Oler AJ, Wollenberg K, Liu Q, Shaffer E, Kozak CA. 2021. Patterns of Coevolutionary Adaptations across Time and Space in Mouse Gammaretroviruses and Three Restrictive Host Factors. Viruses 13:1864.

Boursot P, Din W, Anand R, Darviche D, Dod B, Von Deimling F, Talwar GP, Bonhomme F. 1996. Origin and radiation of the house mouse: mitochondrial DNA phylogeny. Journal of Evolutionary Biology 9:391–415.

Bowles H, Kabiljo R, Jones A, Al Khleifat A, Quinn JP, Dobson RJ, Swanson CM, Al-Chalabi A, Iacoangeli A. 2022. An assessment of bioinformatics tools for the detection of human endogenous retroviral insertions in short-read genome sequencing data. bioRxiv:2022.2002.2018.481042.

Campos-Sánchez R, Cremona MA, Pini A, Chiaromonte F, Makova KD. 2016. Integration and Fixation Preferences of Human and Mouse Endogenous Retroviruses Uncovered with Functional Data Analysis. PLoS Computational Biology 12:e1004956.

Chen X, Li D. 2019. ERVcaller: identifying polymorphic endogenous retrovirus and other transposable element insertions using whole-genome sequencing data. Bioinformatics 35:3913–3922.

Chu C, Borges-Monroy R, Viswanadham VV, Lee S, Li H, Lee EA, Park PJ. 2021. Comprehensive identification of transposable element insertions using multiple sequencing technologies. Nature Communications 12:3836.

Danecek P, Auton A, Abecasis G, Albers CA, Banks E, DePristo MA, Handsaker RE, Lunter G, Marth GT, Sherry ST, et al. 2011. The variant call format and VCFtools. Bioinformatics 27:2156–2158.

Dodding Mark P, Bock M, Yap Melvyn W, Stoye Jonathan P. 2005. Capsid Processing Requirements for Abrogation of Fv1 and Ref1 Restriction. Journal of Virology 79:10571–10577.

Edelmann W, Kröger B, Goller M, Horak I. 1989. A recombination hotspot in the LTR of a mouse retrotransposon identified in an in vitro system. Cell 57:937–946.

Ferraj A, Audano PA, Balachandran P, Czechanski A, Flores JI, Radecki AA, Mosur V, Gordon DS, Walawalkar IA, Eichler EE, et al. 2023. Resolution of structural variation in diverse mouse genomes reveals chromatin remodeling due to transposable elements. Cell Genomics 3.

Frank JA, Singh M, Cullen HB, Kirou RA, Benkaddour-Boumzaouad M, Cortes JL, Garcia Pérez J, Coyne CB, Feschotte C. 2022. Evolution and antiviral activity of a human protein of retroviral origin. Science 378:422–428.

Fujiwara K, Kawai Y, Takada T, Shiroishi T, Saitou N, Suzuki H, Osada N. 2022. Insights into Mus musculus Population Structure across Eurasia Revealed by Whole-Genome Analysis. Genome Biology and Evolution 14.

Fujiwara K, Kubo S, Endo T, Takada T, Shiroishi T, Suzuki H, Osada N. 2024. Inference of selective forces on house mouse genomes during secondary contact in East Asia. Genome Research 34:366–375.

Gardner EJ, Lam VK, Harris DN, Chuang NT, Scott EC, Pittard WS, Mills RE, Consortium TGP, Devine SE. 2017. The Mobile Element Locator Tool (MELT): population-scale mobile element discovery and biology. Genome Research 27:1916–1929.

Goff SP. 2004. Retrovirus Restriction Factors. Molecular Cell 16:849–859.

Huerta-Sanchez E, Jin X, Asan, Bianba Z, Peter BM, Vinckenbosch N, Liang Y, Yi X, He M, Somel M, et al. 2014. Altitude adaptation in Tibetans caused by introgression of Denisovan-like DNA. Nature 512:194–197.

Ikeda H, Sato H, Odaka T. 1981. Mapping of the Fv-4 mouse gene controlling resistance to murine leukemia viruses. International Journal of Cancer 28:237–240.

Inaguma Y, Miyashita N, Moriwaki K, Huai WC, Jin ML, He XQ, Ikeda H. 1991. Acquisition of two endogenous ecotropic murine leukemia viruses in distinct Asian wild mouse populations. Journal of Virology 65:1796–1802.

Kawase M, Ichiyanagi K. 2023. Mouse retrotransposons: sequence structure, evolutionary age, genomic distribution and function. Genes and Genetic Systems 98:337–351.

Keane TM, Goodstadt L, Danecek P, White MA, Wong K, Yalcin B, Heger A, Agam A, Slater G, Goodson M, et al. 2011. Mouse genomic variation and its effect on phenotypes and gene regulation. Nature 477:289–294.

Keane TM, Wong K, Adams DJ. 2012. RetroSeq: transposable element discovery from next-generation sequencing data. Bioinformatics 29:389–390.

Kent TV, Uzunović J, Wright SI. 2017. Coevolution between transposable elements and recombination. Philosophical Transactions of the Royal Society B: Biological Sciences 372:20160458.

Koide T, Moriwaki K, Uchida K, Mita A, Sagai T, Yonekawa H, Katoh H, Miyashita N, Tsuchiya K, Nielsen TJ, et al. 1998. A new inbred strain JF1 established from Japanese fancy mouse carrying the classic piebald allele. Mammalian Genome 9:15–19.

Kojima S, Koyama S, Ka M, Saito Y, Parrish EH, Endo M, Takata S, Mizukoshi M, Hikino K, Takeda A, et al. 2023. Mobile element variation contributes to population-specific genome diversification, gene regulation and disease risk. Nature Genetics 55:939–951.

Lander ES, Linton LM, Birren B, Nusbaum C, Zody MC, Baldwin J, Devon K, Dewar K, Doyle M, FitzHugh W, et al. 2001. Initial sequencing and analysis of the human genome. Nature 409:860–921.

Li H, Durbin R. 2009. Fast and accurate short read alignment with Burrows–Wheeler transform. Bioinformatics 25:1754–1760.

Logsdon GA, Vollger MR, Eichler EE. 2020. Long-read human genome sequencing and its applications. Nature Reviews Genetics 21:597–614.

Mouse Genome Sequencing C, Waterston RH, Lindblad-Toh K, Birney E, Rogers J, Abril JF, Agarwal P, Agarwala R, Ainscough R, Alexandersson M, et al. 2002. Initial sequencing and comparative analysis of the mouse genome. Nature 420:520–562.

Nellåker C, Keane TM, Yalcin B, Wong K, Agam A, Belgard TG, Flint J, Adams DJ, Frankel WN, Ponting CP. 2012. The genomic landscape shaped by selection on transposable elements across 18 mouse strains. Genome Biology 13:R45.

O’Brien SJ, Berman EJ, Estes JD, Gardner MB. 1983. Murine retroviral restriction genes Fv-4 and Akvr-1 are alleles of a single locus. Journal of Virology 47:649–651.

Odaka T, Ikeda H, Moriwaki K, Matsuzawa A, Mizuno M, Kondo K. 1978. Genetic Resistance in Japanese Wild Mice (Mus musculus molossinus) to an NB-Tropic Friend Murine Leukemia Virus2. JNCI: Journal of the National Cancer Institute 61:1301–1306.

Speidel L, Forest M, Shi S, Myers SR. 2019. A method for genome-wide genealogy estimation for thousands of samples. Nature Genetics 51:1321–1329.

Stern AJ, Wilton PR, Nielsen R. 2019. An approximate full-likelihood method for inferring selection and allele frequency trajectories from DNA sequence data. PLoS Genetics 15:e1008384.

Stocking C, Kozak CA. 2008. Endogenous retroviruses. Cellular and Molecular Life Sciences 65:3383–3398.

Storer J, Hubley R, Rosen J, Wheeler TJ, Smit AF. 2021. The Dfam community resource of transposable element families, sequence models, and genome annotations. Mobile DNA 12:2.

Stoye JP, Coffin JM. 1987. The four classes of endogenous murine leukemia virus: structural relationships and potential for recombination. Journal of Virology 61:2659–2669.

Szpiech ZA. 2021. selscan 2.0: scanning for sweeps in unphased data. bioRxiv:2021.2010.2022.465497.

Takada T, Ebata T, Noguchi H, Keane TM, Adams DJ, Narita T, Shin-I T, Fujisawa H, Toyoda A, Abe K, et al. 2013. The ancestor of extant Japanese fancy mice contributed to the mosaic genomes of classical inbred strains. Genome Research 23:1329–1338.

Takada T, Miyazawa H, Yamagata M, Tamura M, Yoshiki A, Toyoda A, Noguchi H, Masuya H. 2025. MoG+3.0: expanded structural variant visualization and integration of genomic data from five newly analyzed mouse strains. Mammalian Genome 37:4.

Tanave A, Koide T. 2020. A role for the rare endogenous retrovirus β4 in development of Japanese fancy mice. Communications Biology 3:53.

Wu T, Yan Y, Kozak Christine A. 2005. Rmcf2, a Xenotropic Provirus in the Asian Mouse Species Mus castaneus, Blocks Infection by Polytropic Mouse Gammaretroviruses. Journal of Virology 79:9677–9684.

Yamada T, Ohtani S, Sakurai T, Tsuji T, Kunieda T, Yanagisawa M. 2006. Reduced Expression of the Endothelin Receptor Type B Gene in Piebald Mice Caused by Insertion of a Retroposon-like Element in Intron 1*. Journal of Biological Chemistry 281:10799–10807.

Zhang Y, Maksakova IA, Gagnier L, van de Lagemaat LN, Mager DL. 2008. Genome-Wide Assessments Reveal Extremely High Levels of Polymorphism of Two Active Families of Mouse Endogenous Retroviral Elements. PLoS Genetics 4:e1000007.

