## Supplementary Figures 1 to 9 for "Evolutionary dynamics of polymorphic endogenous retrovirus insertions across wild house mouse populations"

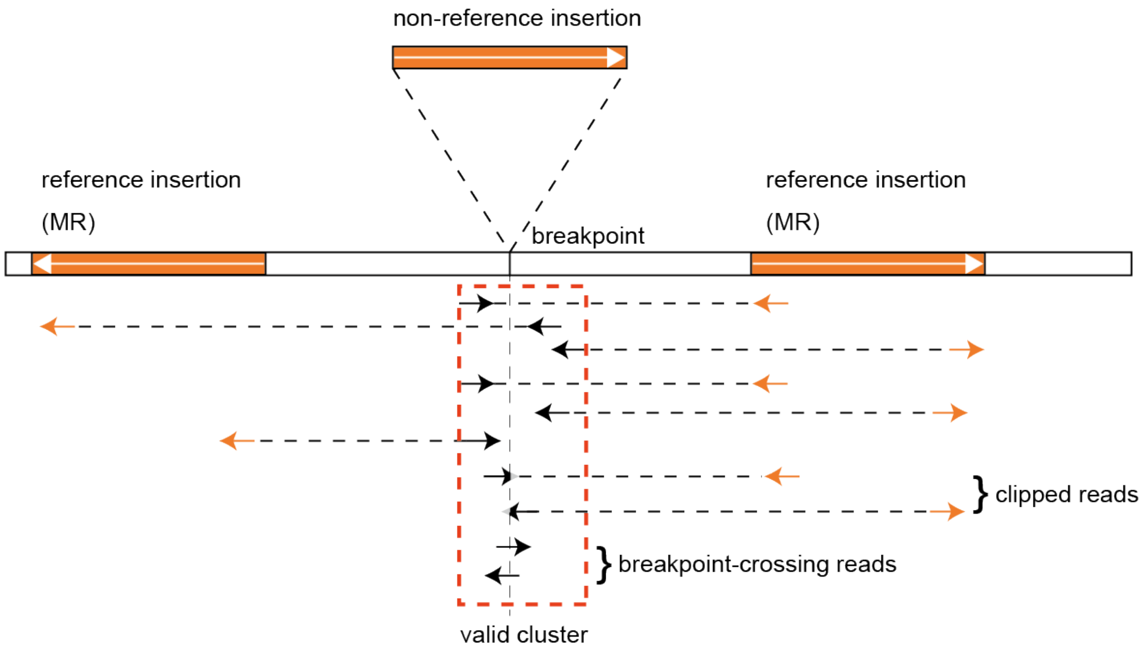


Supplementary Figure 1

Schematic representation of the ERVscanner pipeline. Orange regions represent sequences homologous to ERVs. The topmost bar illustrates a non-reference ERV insertion to be detected. When DNA fragments spanning the insertion site are sequenced, discordant read pairs are generated in which at least one read maps to an ERV-derived masked region (MR) in the reference genome (indicated by orange arrows). The reads indicated by black arrows are referred to as left- and right-supporting reads that define the breakpoint cluster. Breakpoint positions are inferred from clipped reads, whereas breakpoint-crossing reads provide evidence for genotyping.

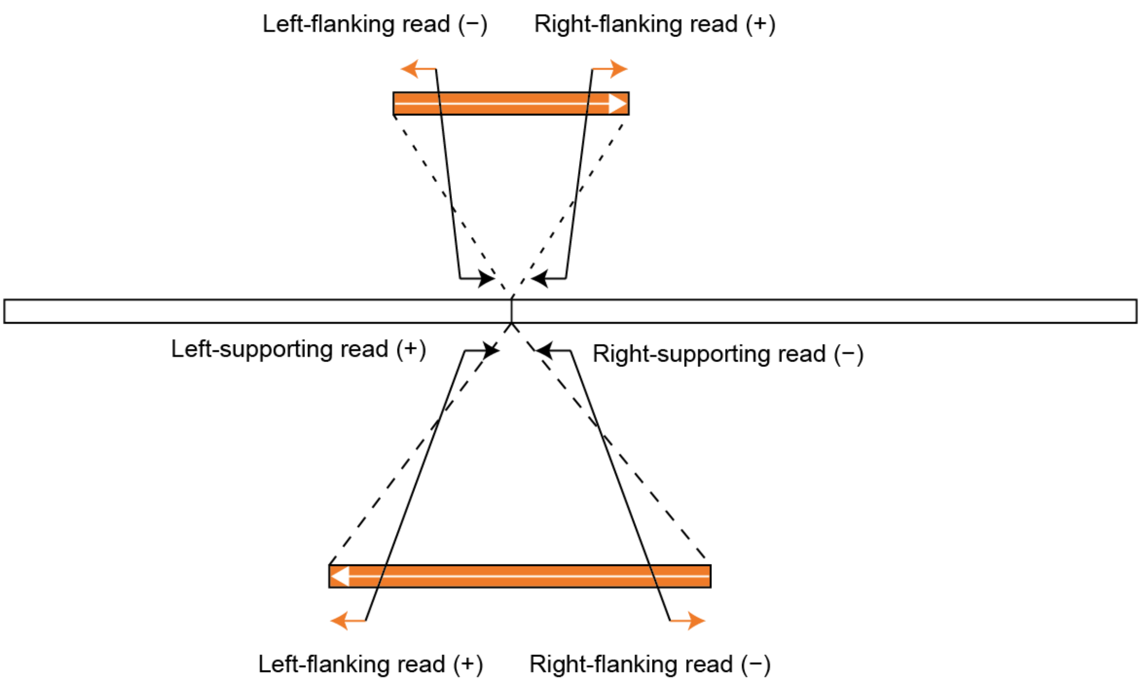


Supplementary Figure 2

Schematic representation of how ERVscanner infers insertion contents. Reads mapped near the insertion site are referred to as supporting reads, and reads paired with the supporting reads are referred to as flanking reads. If the insertion content is correctly inferred, flanking reads from the left and right sides are expected to map to the target sequence with consistent orientations (left − and right +, or left + and right −).

For each target sequence, we counted the minimum number of left- and right-flanking reads with consistent orientations and defined this value as the support score. The target sequence with the highest support score was assigned as the inferred insertion content.

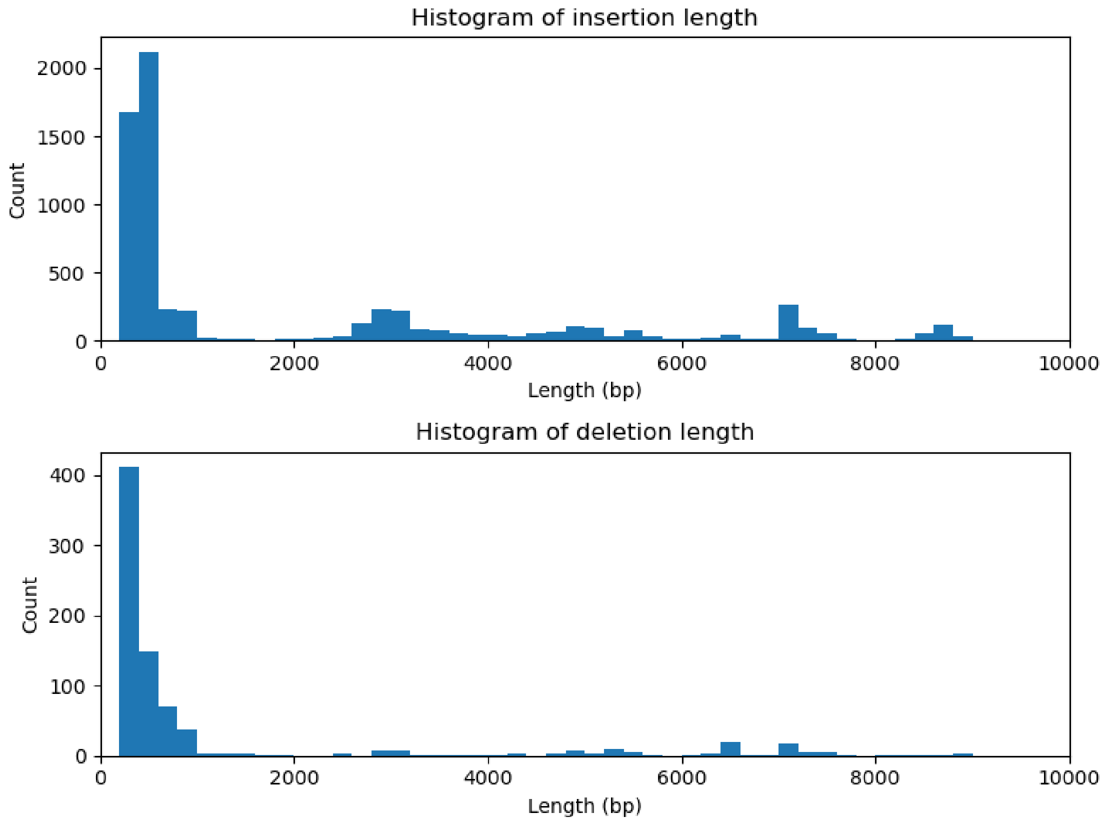


Supplementary Figure 3

Distribution of insertion and deletions length of ERV variants in the JF1 genome, identified using a long-read sequencer.

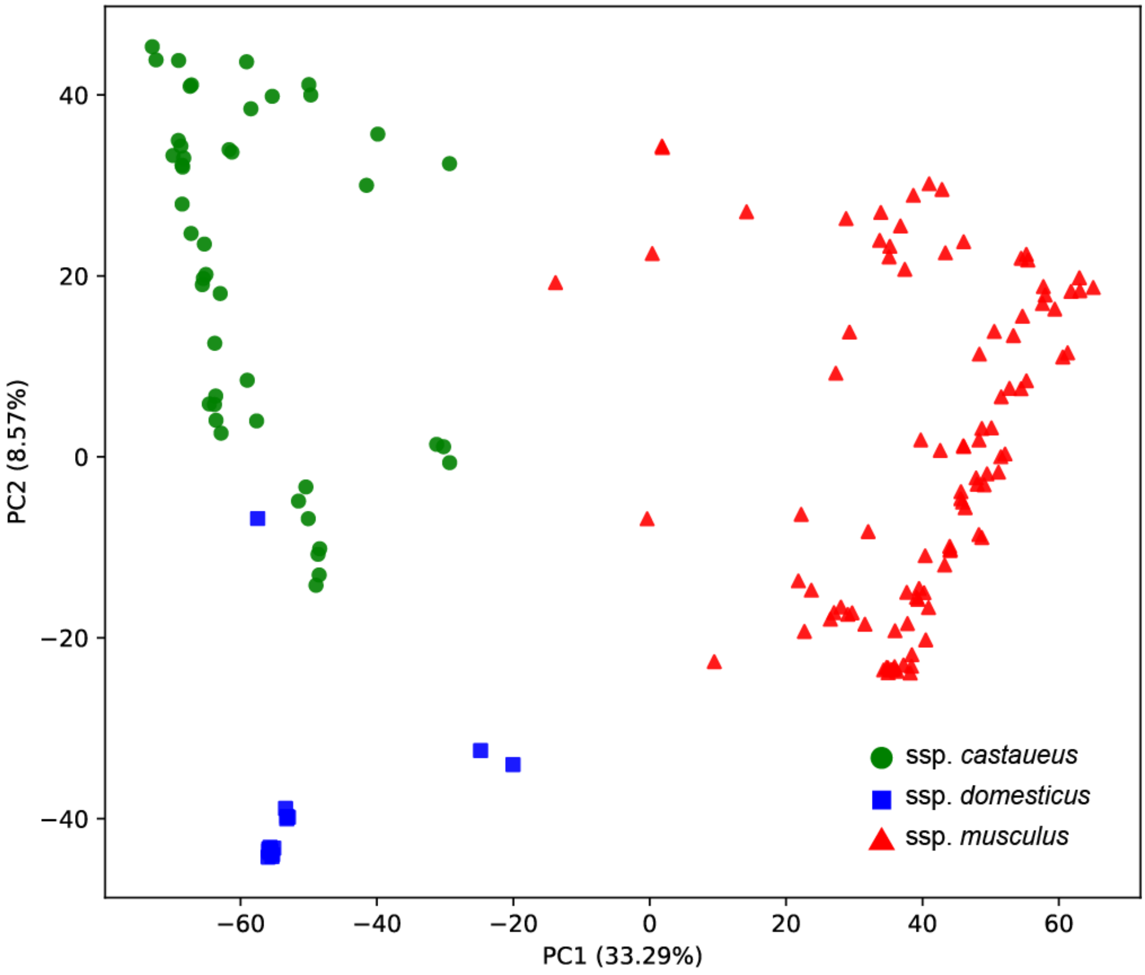


Supplementary Figure 4

PCA plot for 163 wild house mice, using non-reference ERV insertions as genetic markers. PC1 and PC2 were plotted.

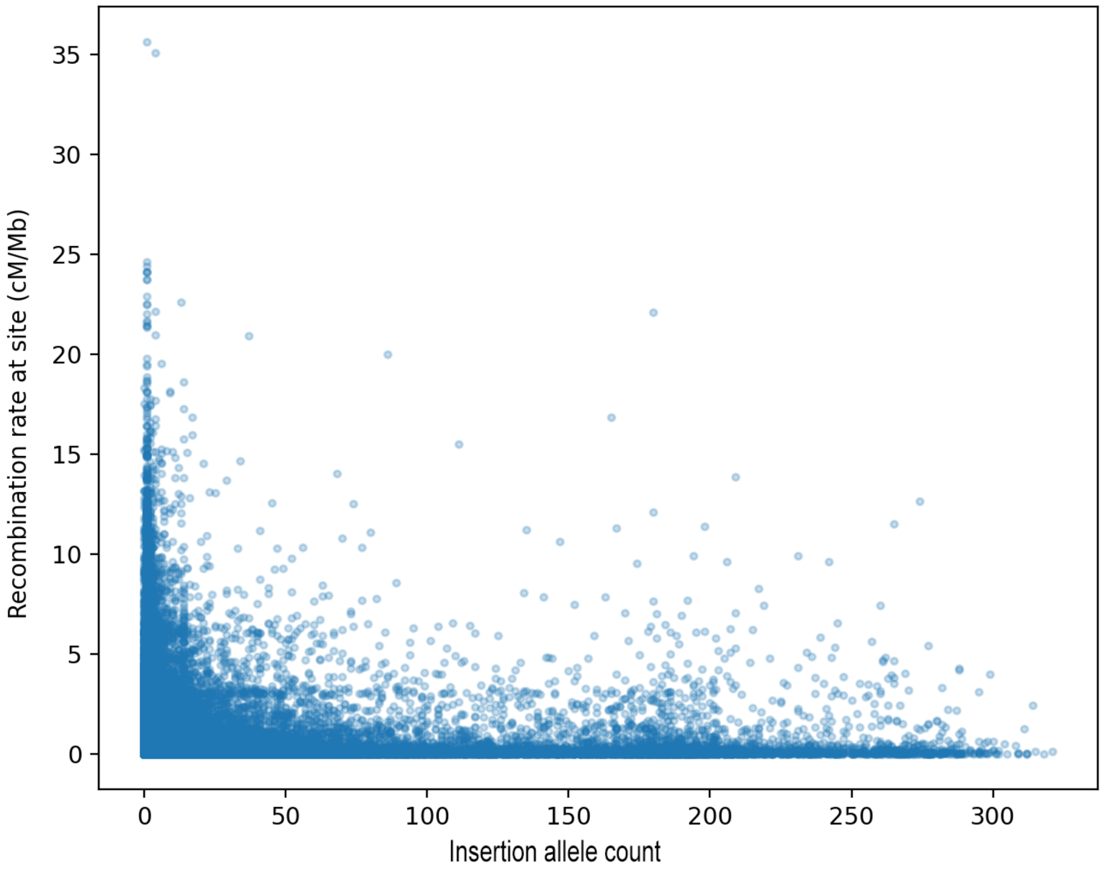


Supplementary Figure 5

Scatter plot of allele count of non-reference insertions of ERVs and recombination rates at site.

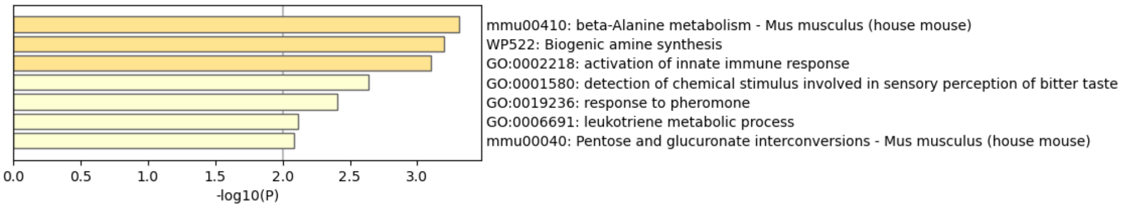


Supplementary Figure 6

Enrichment analysis using Metascape. The enrichment of genes with ERV insertions in their protein-coding sequences is shown.

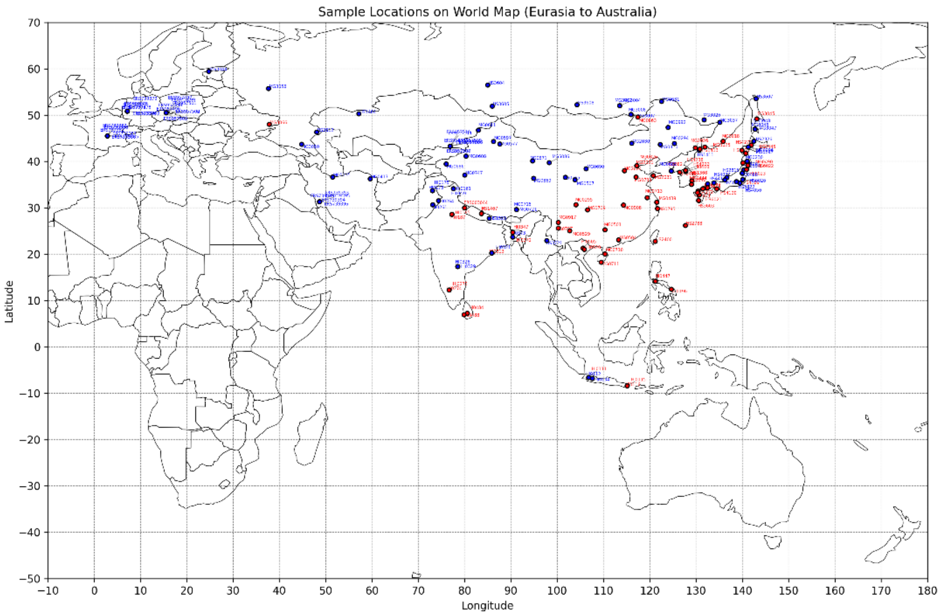


Supplementary Figure 7

Distribution of *Fv4* insertions in Southern and Eastern Eurasia. The blue and red circles represent individuals with the absence and presence of *Fv4*, respectively.

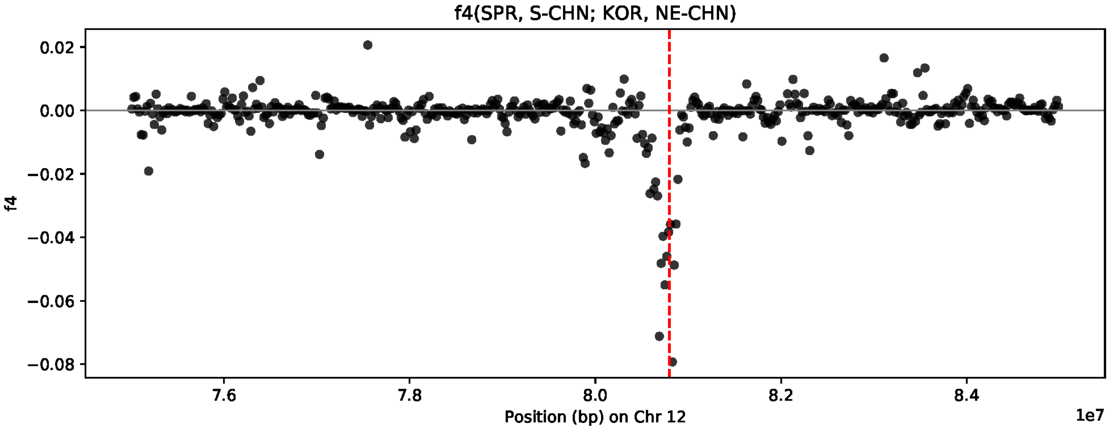


Supplementary Figure 8

$f_4$ statistics were calculated using SNPs in 20-kb windows around the integration site of *Fv4*. The red vertical line represents the inferred integration site of *Fv4*. The $f_4$ statistics were calculated under the configuration $f_4$(SPR, S-CHN; KOR, NE-CHN), where SPR, S-CHN, KOR, and NE-CHN represent *M. spretus*, southern Chinese *M. musculus castaneus*, Korean *M. m. musculus*, and northeastern Chinese *M. m. musculus*, respectively. Assuming that *Fv4* originated from *M. m. castaneus*, negative $f_4$ values indicate introgression from S-CHN into KOR, but not into NE-CHN.

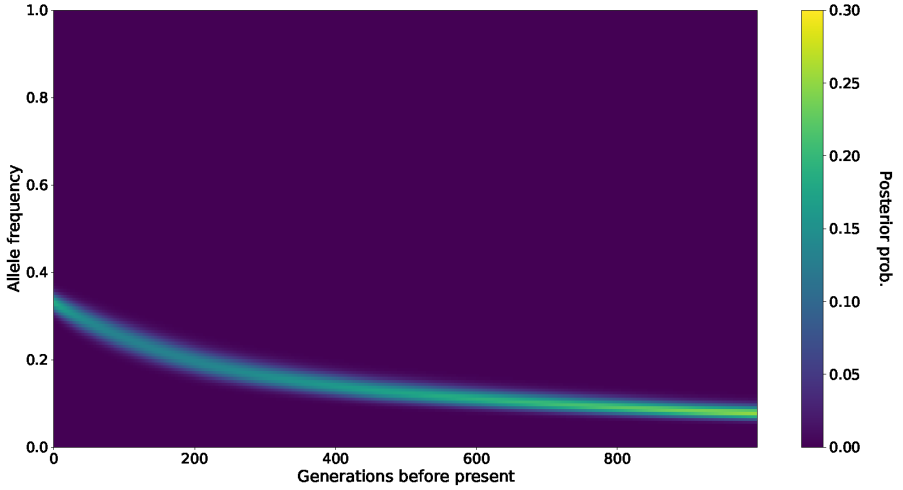


Supplementary Figure 9

Estimated trajectory of Fv4 insertions in the subspecies *musculus*. The genealogy around the *Fv4* insertion was estimated using RELATE, and the trajectory of allele frequency was inferred using CLUES.
